# PyClone-VI: Scalable inference of clonal population structures using whole genome data

**DOI:** 10.1101/2020.08.31.276212

**Authors:** Sierra Gillis, Andrew Roth

## Abstract

We describe PyClone-VI, a computationally efficient Bayesian statistical method for inferring the clonal population structure of cancers. Our proposed method is 10-100x times faster than existing methods, while providing results which are as accurate. We demonstrate the utility of the method by analyzing data from 1717 patients from PCAWG study and 100 patients from the TRACERx study. Software implementing our method is freely available https://github.com/Roth-Lab/pyclone-vi.

## Background

Cancer is an evolutionary process driven by ongoing somatic mutation within the malignant cell population [1, 2]. The combination of mutation, drift, and selection lead to heterogeneity within the population of cancer cells. Identifying population structure and quantifying the amount of heterogeneity in tumours is an important problem which has been extensively studied [3, 4, 5, 6, 7, 8]. High throughput sequencing (HTS) provides a powerful approach to solve the problem with both bulk and single cell approaches being employed. While single cell sequencing approaches can more accurately resolve clonal population structure, they are not widely available and have limitations both technical and due to cost. Using bulk sequencing to study heterogeneity thus remains the predominate approach, and methods for studying heterogeneity using bulk sequencing will become even more important as HTS is increasingly used in translational and clinical work [9, 10, 11, 12].

Identifying population structure and quantifying heterogeneity from bulk sequencing data is a computationally challenging problem. The core issue is to deconvolve sequence data generated from a mixture of cell populations. This task is challenging because neither the genotypes of the populations nor the number of populations is known. In addition, factors such as tumour cellularity and copy number variation co-incident to small nucleotide variants (SNVs) further complicate the analysis.

The past decade has seen a number of methods to deconvolve bulk data and infer clonal population structure, in particular to identify populations using SNV data. One of the first approaches developed was PyClone, which remains widely used. PyClone was originally developed for use with small panels of deeply sequenced mutations as input [4]. While the PyClone method can in principle be applied to genome scale analysis, the computational cost becomes prohibitive. This deficiency has limited the utility of PyClone for the analysis of genome scale datasets with 10,000s – 100,000s of mutations. In this work we present a new tool, which we refer to as PyClone-VI, which is orders of magnitudes faster than the original PyClone method, while providing comparable accuracy.

### Related work

A number of other methods have been developed to efficiently infer clonal population structure from genome scale data. We provide a brief, non-extensive, review of some of the most popular methods.

SciClone uses Bayesian mixture models and variational inference (VI) like our proposed approach PyClone-VI [6]. However, because SciClone fails to correct for coincident copy number variation, it is only applicable to clustering mutations in regions with no copy number variation or with single copy deletions. It follows that in practice SciClone cannot be applied to many tumours, especially when multi-region sequencing is performed, as few mutations will fall in such regions.

EXPANDS is based on the principle of clustering probability distributions of cancer cell fractions (CCFs) using a multi-stage optimization procedure [5]. It has been applied to whole genome studies alongside PyClone and shown to perform similarly [13]. One key difference between EXPANDS and PyClone is that mutations are clustered independently in each sample and then the clusters are combined in a post-processing step. As a result of post-hoc analysis, statistical strength cannot be shared between samples when inferring population structure using EXPANDS. QuantumClone is a Bayesian mixture model that is fit to the data using expectation maximization (EM) to find the maximum a posteriori (MAP) estimate [8]. MAP estimation for mixture models is prone to overfitting, in the sense that the model will tend to use all possible clusters (clones). To address the model selection problem QuantumClone uses the Bayesian Information Criterion (BIC) to select the number of clusters. QuantumClone can correct for genotype effects and jointly analyse multi-region data. The use of the BIC for model selection requires that multiple runs of the method be performed with varying numbers of clusters. QuantumClone is conceptually similar to our proposed method, however our approach avoids the expensive model complexity search across varying number of clusters. As we demonstrate in the experiments, avoiding restarts for the model complexity search can lead to a considerable reduction in runtime.

PhyloWGS is a popular approach which attempts to solve a more challenging problem of identifying not only clonal populations, but the phylogeny that relates them [7]. PhyloWGS adopts a very similar model to PyClone, but substitutes the Dirichlet process prior for clustering with a tree structured stick breaking prior [14]. Like PyClone, PhyloWGS relies on Markov Chain Monte Carlo (MCMC) methods and can be computationally expensive to run with large datasets.

## Results

### PyClone-VI is as accurate as PyClone but faster

PyClone-VI introduces two levels of approximation to the original PyClone model. First, we alter the model to make it more tractable to perform variational inference. Second, we use variational inference which is an approximate method to infer a posterior distribution. To assess the impact these approximations have and investigate whether they lead to tangible performance gains, we compared PyClone-VI to PyClone using synthetic data. We simulated data from the PyClone model with varying numbers of mutations. We generated datasets with 50, 100 and 1000 mutations. Each simulated dataset had four samples each with a tumour content of 1.0. Total copy number for each loci ranged from one to four and major copy number was allowed to vary from one to the total copy number. Genotypes were simulated by selecting whether mutations were late events which affected only one copy or early events which occurred on either the major or minor allele before the copy number change. We simulated the depth of coverage from a Poisson distribution with mean 100. We repeated the simulation for each number of mutations 100 times to generate 300 datasets in total.

The results of this analysis are summarized in Figure 1. Clustering accuracy was assessed using the V-Measure metric with a value of 1.0 indicating perfect accuracy (Figure 1 **a**) [15]. The mean difference in V-Measure between PyClone and PyClone-VI was 0.011 in favour of PyClone. To assess the accuracy of the cancer cell fraction (CCFs) estimates we computed the mean absolute deviation of the predicted CCF from truth for each mutation (Figure 1 **b**). The mean difference in CCF error was 0.00036 in favour of PyClone-VI. These results suggest there is a negligible performance difference between the two approaches. We note that we would expect PyClone to have a slight performance advantage in this experiment as we simulated the data from the PyClone model rather than the PyClone-VI model. Finally, we sought to quantify the computational performance of both methods. Figure 1 **c**) and **d**) show the runtime and maximum memory used by both methods. PyClone-VI outperforms PyClone in terms of runtime by nearly two orders of magnitude regardless of the number of mutations (Figure 1 **c**). PyClone-VI also uses significantly less memory than PyClone (Figure 1 **d**). Theoretical memory usage for the original PyClone method scales as *𝒪*(*n*^2^) where *n* is the number of mutations. In contrast, memory usage for PyClone-VI scales as *𝒪* (*n*). The empirical results in Figure 1 **d** appear to support this.

**Figure 1.**
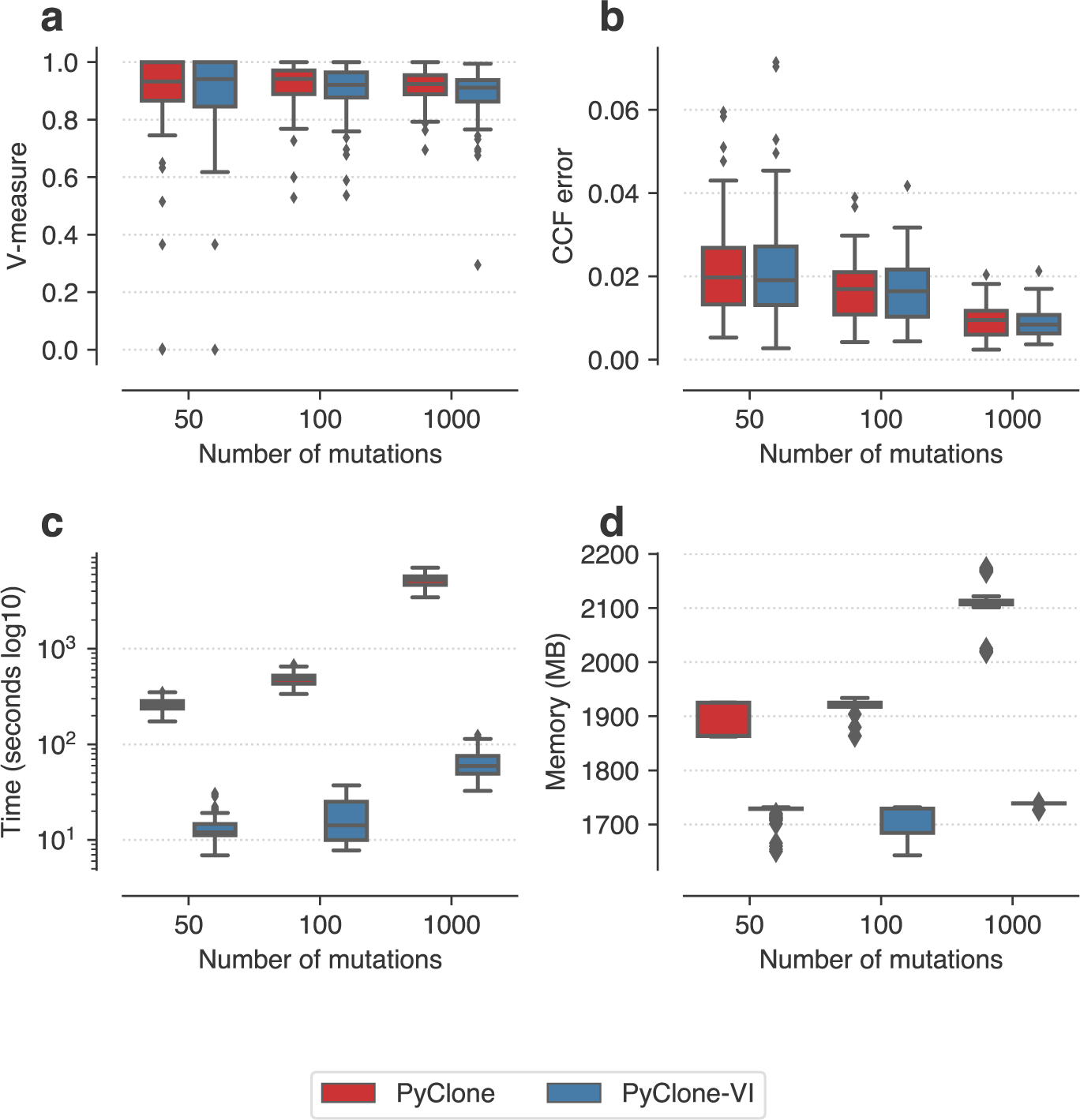
Comparison of PyClone and PyClone-VI. **a**) V-measure as a function of the number of mutations. **b**) Mean absolute deviation of inferred CCF from truth as a function of the number of mutations. **c**) Runtime of the methods. **d**) Memory usage.

### PyClone-VI is significantly faster than existing methods

We next sought to compare the performance of PyClone-VI against other state of the art methods. In addition to comparing against PyClone, we also considered PhyloWGS and QuantumClone. We downloaded synthetic data used in the ICGC-TCGA DREAM Somatic Mutation Calling – Tumour Heterogeneity Challenge, an open competition to benchmark methods for studying clonal heterogeneity [16]. We limited the analysis to tumours with 10,000 mutations or fewer due to issues relating to runtime (PyClone, PhyloWGS and QuantumClone) and memory (PyClone and QuantumClone). As in the previous experiment, we consider two metrics to assess performance: V-measure (Figure 2 **a**) and mean absolute deviation error in predicted CFF per mutation (Figure 2 **b**).

**Figure 2.**
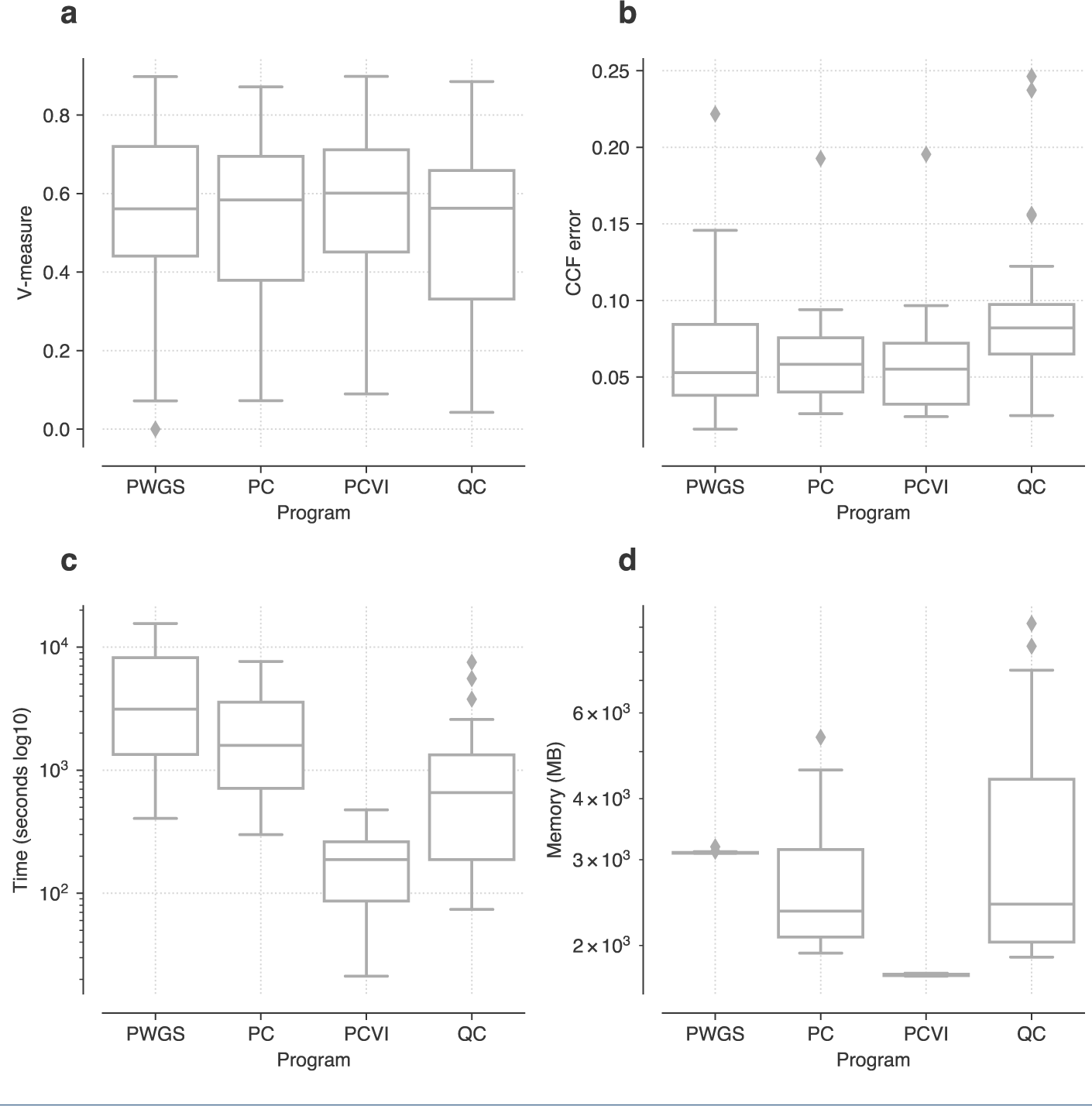
Analysis of the DREAM SMC-Het data. Analysis of the CGC-TCGA DREAM Somatic Mutation Calling – Tumour Heterogeneity Challenge data using PhyloWGS (PWGS), PyClone (PC), PyClone-VI (PCVI) and QuantumClone (QC). **a**) Comparison of V-measure across the methods (higher is better). **b**) Comparison of the mean absolute deviation of estimated cancer cell fraction across methods (lower is better). **c**) Comparison of runtime across methods (lower is better). **c**) Comparison of memory usage across methods (lower is better).

When comparing methods we applied the Friedman test to see if there were any significant differences in performance between the methods (p-value *<* 0.01). If the Friedman test was significant we then applied the post-hoc Nemenyi test with a Bonferroni correction to all pairs of methods to determine which methods showed significantly different performance from each other (p-value *<* 0.01) [17]. All statements of significance are with respect to this test.

PyClone-VI significantly outperformed PyClone and QuantumClone with respect to clustering performance. Though PyClone-VI performed better on average than PhyloWGS the difference was not significant (*p* = 0.46). With respect to accuracy estimating CCF, both PyClone-VI and PhyloWGS outperformed QuantumClone. There were no other significant differences in accuracy metrics between methods.

In general, the results were quite similar across methods, with the differences in performance being quite small. However, there was a significant difference in runtime between methods. PyClone-VI was significantly faster and more memory efficient than all other approaches, finishing 10x-100x times faster than the other approaches while requiring less memory (Figures 2 **c** and **d**). A caveat to this analysis is that runtime is a tuneable parameter for all these approaches. Fewer MCMC iterations can be performed for PyClone and PhyloWGS to shorten runtime at the expense of accuracy. Similarly, QuantumClone and PyCloneVI can use fewer random restarts to speed up runtime, again trading accuracy. For this analysis we attempted to select parameters which gave comparable accuracy (see methods). We did not make use of parallel computing in this experiment. Both QuantumClone and PyClone-VI can perform random restarts in parallel to decrease runtime. The MCMC based methods cannot be parallelised in the same way.

### Analysis of PCAWG cohort

To demonstrate the real life utility of PyClone-VI we analysed the data from the Pan-Cancer Analysis of Whole Genomes (PCAWG) [18]. We downloaded processed data from the ICGC data portal and pre-processed it for input into PyClone-VI. The only filtering performed was to remove mutations with no copy number information or in regions with total copy number zero. We analysed the resulting data from 1717 patients with 28 to 881464 mutations. All data was single sample whole genome data. Figure 3 **a**) shows the runtime of PyClone-VI as function of the number of mutations. Runtime increases linearly with the number of mutations with times ranging from 11-28575 seconds. Figure 3 **b**) shows the runtime as a function of the number of clones detected and Figure 3 **c**) shows how the number of clones detected depends on the number of mutations. The trend is that more clones are detected as more mutations are included, with runtime correspondingly increasing with the number of clones. Figure 3 **d**) is an illustrative analysis which shows the number of clones normalised by the number of mutations broken down by ICGC project.

**Figure 3.**
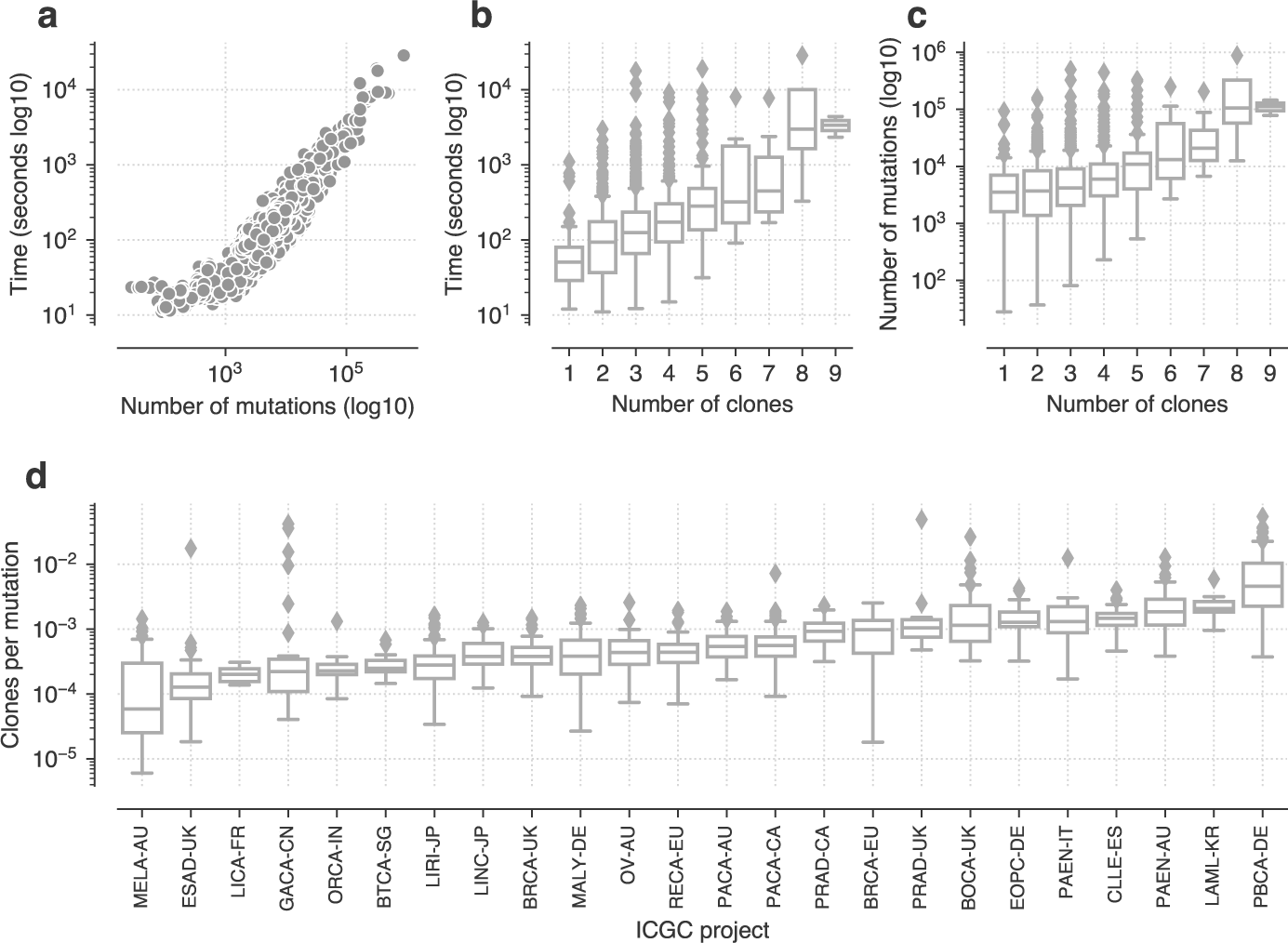
Analysis of the PCAWG cohort. **a**) Runtime of PyClone-VI as a function of the number of mutations. **b**) Runtime of PyClone-VI as a function of the number of clones inferred. **c**) Comparison between the number of clones found and number of mutations. **d**) Number of clones normalized by total number of mutations for each ICGC project.

### Analysis of TRACERx cohort

As another real world demonstration, this time with multiple samples, we analysed whole exome data from the 100 lung cancer patients from the TRACERx study [12]. Patients had between 1-7 samples sequenced from different regions of their tumours with between 65 and 3566 mutations detected. Figure 4 **a**) shows the runtime of PyClone-VI as function of the number of mutations. Again runtime increases linearly with the number of mutations with times ranging from 9-1454 seconds. Figure 4 **b**) and **c**) show runtime and runtime normalised by the number of mutations with varying numbers of samples. Runtime does not directly increase with the number of samples (Figure 4 **b**), but once the runtime is normalised to account for the number of mutations we see an increase (Figure 4 **c**). In Figure 4 **d**) and **e**) we show the number of mutations and clones that can be resolved as a function of the number of samples. Interestingly, the number of mutations identified does not seem to depend strongly on the number of samples, however the number of clones which can be detected increases as more samples are added. This result illustrates the important role that multi-region sequencing plays in determining clonal population structure. Eight patients in the cohort had only a single sample. We compared the number of mutations in these patients inferred to be clonal to the number inferred to be clonal from multi-region sequencing (Figure 4 **f**). The proportion of detected clonal mutations decreases in the multi-sample setting suggesting that many apparently clonal mutations in single sample sequencing may in fact be subclonal, consistent with the findings in [12] which performed a more thorough held out sampling.

**Figure 4.**
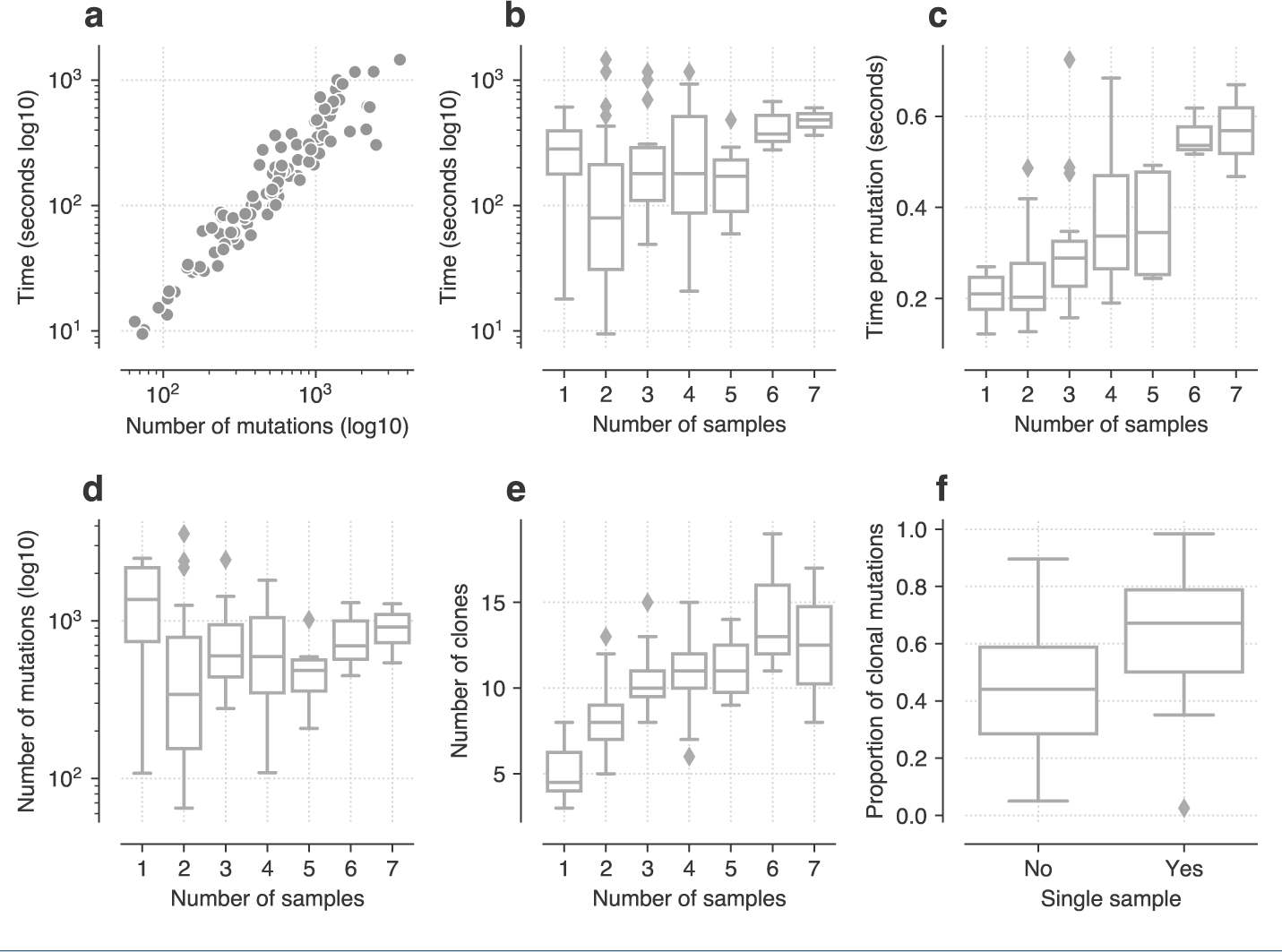
Analysis of the TRACERx cohort. **a**) Runtime of PyClone-VI a function of the number of mutations. **b**) Runtime of PyClone-VI a function of the number of samples. **c**) Runtime normalised by number of mutations for varying numbers of samples. **d**) Number of mutations detected with varying numbers of samples. **e**) Number of clones detected with varying numbers of samples. **f**) Comparison of proportion of mutations deemed clonal when using single versus multiple samples.

## Discussion

PyClone-VI achieves significant computational gains over the original PyClone method by altering the model and changing the approach used for inference. To do so we introduce several approximations on top of those already in the PyClone model.

We assume that CCF values can only take on a finite set of values. The number of possible values determines the accuracy of this approximation and the runtime. For the analyses performed in this paper we used a grid of 100 values, which provides CCFs accurate to within 0.01. Using a larger grid of values will provide more accurate estimates if the mutations are sequenced to a sufficient depth. In general, large numbers of mutations are not deeply sequenced, so using relatively sparse grids is appropriate for the data. If a small panel of mutations is deeply sequenced, then the original PyClone method maybe more appropriate than PyClone-VI.

Another approximation we make is to use a finite mixture model in place of a Dirichlet process (DP) for clustering. We rely on the variational inference procedure to automatically perform model selection by only using the number of clusters supported by the data. The approach of using more clusters than needed is heuristic, however it is widely employed and generally performs well [19]. We note neither DP models or using the BIC are guaranteed to consistently estimate the correct number of clusters.

The use of VI rather than MCMC for inference means that PyClone-VI will deliver posterior approximations of unknown accuracy. In contrast, MCMC approaches are guaranteed to approximate the posterior to arbitrary accuracy given enough samples are drawn. In practice, VI approaches are typically observed to estimate the mean of the posterior distribution well, but to underestimate the variance. When inferring clonal population structure the underestimation of variance would lead to over confident assignment of mutations to clusters and under-estimates of error bar widths for CCF values. If accurate estimates of these values are required, then we recommend the use of the original PyClone model. It is our observation that most users do not make use of these values, and instead rely on the point estimates generated by PyClone. In this case, PyClone-VI should be the preferred approach due to reduced runtime.

Like PyClone, PyClone-VI clusters mutations which share the same evolutionary history. Such mutations originate at the same point in the phylogeny and exhibit the same pattern of mutation loss. PyClone-VI does not attempt to infer the phylogenetic tree, in contrast to methods such as PhyloWGS. Ignoring the phylogenetic structure is a potential weakness, but it does mean we do not have to make additional assumptions such as mutations cannot be lost once gained. Such assumptions are restrictive and violated in many cancers [20]. We believe that the ability to quickly cluster mutations will be useful for downstream software which attempts to infer phylogenies. By reducing the size of the input data from the number of mutations to the number of clonal populations, more sophisticated and computationally expensive tree building methods can be used [21, 22, 23].

## Conclusions

We have introduced a new method, PyClone-VI, for inferring clonal population structure in tumours from point mutations measured using high throughput sequencing. PyClone-VI is significantly more computationally efficient than existing approaches and provides comparable accuracy. Tumours with 100,000s of mutations can easily be analysed by PyClone-VI in less than a day on a personal computer, a dramatic reduction in both runtime and memory required for this analysis. PyClone-VI will be a useful tool for researchers performing large cohort studies of tumour heterogeneity. PyClone-VI will also be useful in clinical studies which integrate WGS analysis of tumours and require timely analysis to inform treatment decisions.

## Methods

Inference in the original PyClone package was performed using Markov Chain Monte Carlo (MCMC) sampling [4]. As the number of mutations grows, each iteration of the MCMC sampler becomes slower which is problematic as large datasets likely need many more iterations of MCMC sampling than small datasets which further adds to the computational complexity. However, many users do not adjust for this factor, and as result PyClone is often run with too few iterations for the MCMC chain to converge leading to poor performance. One widely observed symptom of this problem is the tendency for PyClone to produce many clusters containing a single mutation [8].

To overcome these limitations we have modified the original PyClone model. This modification has allowed us to develop and implement an efficient VI procedure which is orders of magnitudes faster than the previous MCMC method. We refer to this new model and software implementation as PyClone-VI. In addition to being significantly faster, this approach also removes the need for the user to assess the convergence of the MCMC sampler thus reducing potential for misuse.

### PyClone

We provide a brief review of the original PyClone method here to motivate the changes in Pyclone-VI. More details can be found in the original PyClone paper [4] which includes additional details such as how to elicit genotype priors and the form of the emission distributions supported.

The original PyClone model is a Dirichlet process (DP) mixture model [24]. The basic hierarchical model is as follows

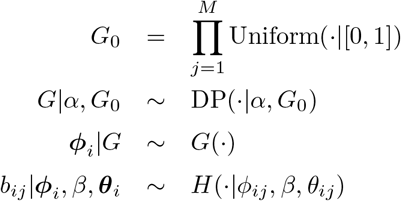

Here we use the distribution *H* to denote the emission distribution used to generate the observed variant read counts *b*_*ij*_, where *i* indexes the mutation and *j* the sample. This distribution depends on local hyper-parameters *θ*_*ij*_ which capture information about the genotype and read depth. The parameter *β* represents global hyper-parameters which are shared across mutations. In the original PyClone paper when using a Beta-Binomial distribution *β* would be the precision of the distribution.

The above model induces a clustering of mutations since the measure *G* sampled from the DP is almost surely discrete which implies there is a non-zero probability that mutations share the same CCF. We can define a clustering of the mutations as follows, let 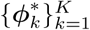 be the unique set of CCFs used to generate the data. Then for mutation *i* we define *z*_*i*_ = *k* if 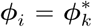. The introduction of the cluster indicator variable *z*_*i*_ is commonly used when developing MCMC sampling strategies for DP mixture models [25]. This formulation is also useful for allowing us discuss how to modify the PyClone model to derive a more computationally efficient approach.

The original PyClone model makes use of the DP to solve the model selection problem. The model selection problem refers to the fact we do not know the true number of clusters (clones) in the model. The DP formulation solves this by positing there exists an infinite number of clusters, but the observed data will only be generated from a finite subset of these. While DP mixtures provide an elegant solution to the model selection problem, they tend to be computationally expensive. The computational expense primarily due to the need to use MCMC methods to approximate the posterior distribution and thus infer model parameters [25].

### Variational inference

VI is a popular alternative to MCMC methods in the Bayesian statistics and machine learning literature [26]. VI reformulates the problem of approximating the posterior as an optimization problem. In the general case, a variational distribution *q*(*θ*|*λ*) is assumed, where *θ* are the model parameters and *λ* are the variational parameters. The goal is to find the variational distribution *q*(*θ*|*λ*) that minimizes some notion of distance from the posterior distribution *p*(*θ*|*X*). A widely used measure of distance is the *exclusive* Kullback-Leibler divergence denoted KL(*q*|*p*).

VI using KL(*q*|*p*) as the objective can lead to efficient inference procedures that provide adequate approximations to the true posterior for many problems. Mean field VI (MFVI) often called variational Bayes in the machine learning and statistics literature posits the variational distribution decomposes as a product of terms for each model parameter *q*(*θ*|*λ*) = ∏_*s*_ *q*(*θ*_*s*_|*λ*_*s*_). For models which obey certain conjugacy constraints, simple closed form MFVI updates can be derived leading to efficient inference algorithms. The updates take the form

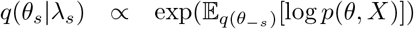

where 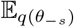 denotes the expectation taken over all parameters except *θ*_*s*_ [27]. The need to compute an expectation is what leads to the constraints on conjugacy for MFVI. We note there has been significant work recently on using Monte Carlo methods to compute these expectations in models that don’t satisfy conjugacy constraints [28, 29]. These approaches could potentially be used as an alternative to our proposed method for performing VI for the PyClone model.

The original PyClone model does not fall in the class of models for which MFVI is easily applicable. There are two issues. The most important issue is the emission density *H* does not have a conjugate prior distribution. The second related issue is that while there are ways to perform VI with DP mixtures, they require that we have a conjugate emission density [30]. Moreover these approaches impose a finite truncation on the number of clusters. This latter point means there is not a major advantage to using the DP when employing VI [31]. Rather, using over complete finite mixture models is often equally effective. Here we use over complete to mean we fit a finite mixture model with more components than we expect to need [32], and allow the inference procedure to perform model selection [19].

### PyClone-VI model

In order to apply VI to fit the PyClone model, we make some modifications to the model. First, we change the model from a DP mixture model to a finite mixture model. In principle the use of a finite mixture model means we must address the model selection problem and fit the model with a varying number of clusters *K*. In practice we avoid this issue by setting *K* to be large and allowing the inference procedure to only use the number of clusters required. This heuristic strategy has been shown to work well in practice [19, 33]. The second modification is to assume that the CCFs of mutations *φ*_*ij*_ can only take values in a finite set 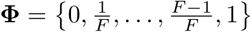 where |**Φ**| = *F* + 1. This change is primarily motivated by computational considerations, but can be justified by noting that we typically sequence genomes to 50x – 1000x when performing whole genome or exome sequencing. Thus, it would seem unreasonable to expect to resolve the CCF of a mutation to arbitrary precision. Provided we choose the grid of CCF values to be sufficiently large, this approximation should thus yield reasonable results.

The modified version of the PyClone model which we call PyClone-VI is defined as follows

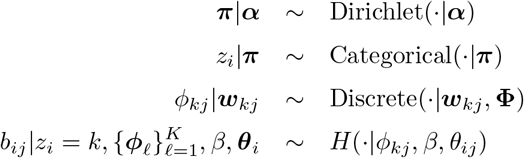

where Discrete(*·*|***w*, Φ**) indicates the discrete distribution with mass vector ***w*** and support **Φ**. We use the uninformative priors ***α*** = *α*1_*K*_, where 1_*K*_ is the vector of ones of length *K*, and 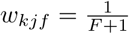.

The joint distribution is thus given by

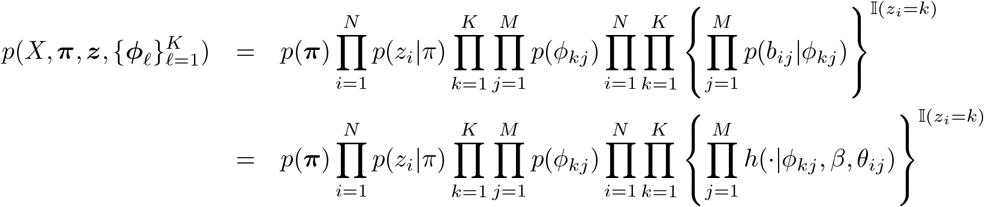

where we have suppressed the dependence on hyper-parameters for notational clarity. We let *h*(*·*|*φ*_*kj*_, *β, θ*_*ij*_) denote the emission density and 𝕀(*z*_*i*_ = *k*) the indicator function which is one when *z*_*i*_ = *k* and zero otherwise. As we will show in the next section this formulation leads to an efficient MFVI procedure.

### Inference

We use MFVI to fit the PyClone-VI model. To do so we make the usual mean field assumption for our variational distribution *q*.

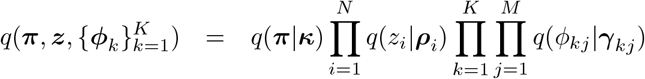

The distributional assumptions are as follows

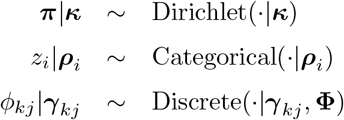

The densities are then given by

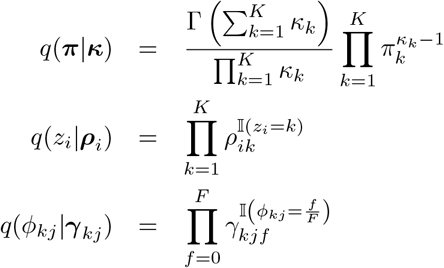

Thus we need to optimize the variational parameters 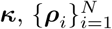 and 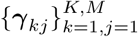. The parameter updates can be derived by applying the standard MFVI update.

Thus we have

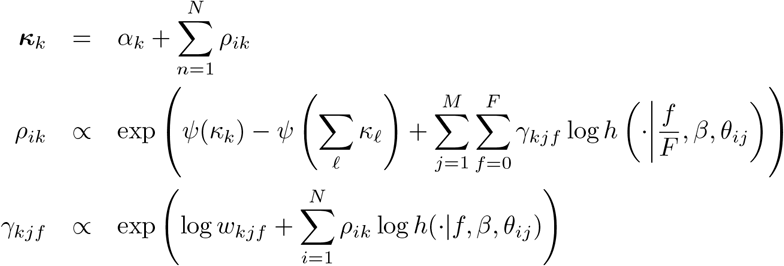

and we have the following normalization constraints

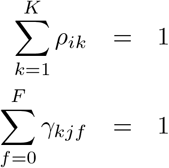

These updates are iterated until convergence. Convergence can monitored by computing the difference in the evidence lower bound (ELBO) after each update [26]. Monitoring the ELBO is also useful to assess that the software implementation is correct, as it should increase monitonically.

Since we assume the CCFs, *φ*_*ij*_, can only take a finite set of values we can evaluate *h*(*·*|*φ*_*kj*_, *β, θ*_*ij*_) for all mutations and samples across this grid as a pre-processing step during inference. Caching this value leads to a dramatic reduction in runtime for the method. This strategy is only applicable if the global parameters *β* of the emission distribution *h* are fixed. In practice, this means we fix the precision term of the Beta-Binomial emission distribution, rather than estimating it as PyClone does. We also treat the hyper-parameter *α* as a fixed parameter. This hyper-parameter weakly controls the number of clusters used, with values greater than one promoting the use of more clusters, and values less than one fewer. For all experiments in this work we used a value of one.

### Experiments

#### Synthetic data

We simulated data from the PyClone model with 50, 100 and 1000 mutations using a DP concentration parameter of 1.0. Additional simulation parameters are described in the results. We used PyClone version 0.13.1 run with 10,000 iterations and discarding the first 1,000 as burn-in. We ran PyClone-VI using 40 clusters and 100 random restarts.

#### DREAM data

We downloaded the ICGC-TCGA DREAM Somatic Mutation Calling – Tumour Heterogeneity Challenge from www.synapse.org. A custom script was used to process the battenberg TSV and mutect VCF files for input into PyClone, PyClone-VI and QuantumClone. We used the included PhyloWGS parser for these input formats to generate input files for PhyloWGS. Tumour content values were set to the ground truth values provided for all methods which accept this argument. PhyloWGS was run for 10 iterations of burn-in and subsequently 100 samples were collected from the MCMC trace. We selected the maximum a posteriori sample, that is the sample with the highest joint probability, to compute estimates from PhyloWGS. PyClone was run for 1,000 iterations, discarding the first 100 iterations as burn-in. We used the PyClone Beta-Binomial emission distribution with the *connected* initialization strategy and *major copy number* prior elicitation method. Default parameters were used for post-processing the PyClone MCMC trace. QuantumClone was run with 2-10 clones and 10 random restarts. PyClone-VI was run with 10 clusters, 100 random restarts and used the Beta-Binomial emission distribution.

#### PCAWG data

We downloaded SNV and CNV data from PCAWG project hosted in the ICGC portal [18]. We used a custom script to pre-process the data into a format compatible with PyClone-VI, extracting read counts from the input VCF files and allele specific copy number from the CNV data. We ignored sub-clonal CNVs and removed mutations with major copy number zero. We fit PyClone-VI using the Binomial emission distribution with 20 clusters and 100 random restarts.

#### TRACERx data

We downloaded SNV and CNV data included in the supplementary material of [12]. We used a custom script to pre-process the data into a format compatible with PyClone-VI. We fit PyClone-VI using the Binomial emission distribution with 40 clusters and 100 random restarts.

## Author’s contributions

AR conceived and implemented the method, performed the experiments and wrote the text. SG helped with performing experiments and writing the text.

## Acknowledgements

We would like to thank Hoa Tran for her feedback during manuscript preparation. We would like to thank Alexandre Bouchard-Côté for helpful discussion about how to develop the approximate inference procedure.

## Additional Files

Additional file 1 — Supplementary tables

Excel file containing supplementary tables supporting figures and statistical results in the paper.

- S1 – Performance results for the comparison of PyClone and PyClone-VI using synthetic data used in Figure 1
- S2 – Performance results for the analysis of DREAM SMC-HET data used in Figure 2
- S3 – Friedman test results for comparing methods using the DREAM SMC-HET data.
- S4 – Post-hoc Nemenyi test for comparing methods using the DREAM SMC-HET data.
- S5 – Results from the PCAWG data analysis used in Figure 3.
- S6 – Results from the TRACERx data analysis used in Figure 4.

